# Prolonging the integrated stress response enhances CNS remyelination in an inflammatory environment

**DOI:** 10.1101/2020.12.29.424699

**Authors:** Yanan Chen, Rejani B Kunjamma, Molly Weiner, Jonah R. Chan, Brian Popko

## Abstract

The inflammatory environment of demyelinated lesions in multiple sclerosis (MS) patients contributes to remyelination failure. Inflammation activates a cytoprotective pathway, the integrated stress response (ISR), but it remains unclear whether enhancing the ISR can improve remyelination in an inflammatory environment. To examine this possibility, the remyelination stage of experimental autoimmune encephalomyelitis (EAE), as well as a mouse model that incorporates cuprizone-induced demyelination along with CNS delivery of the proinflammatory cytokine IFN-γ were used here. We demonstrate that either genetic or pharmacological ISR enhancement significantly increased the number of remyelinating oligodendrocytes and remyelinated axons in the inflammatory lesions. Moreover, the combined treatment of Sephin1 with the oligodendrocyte differentiation enhancing reagent bazedoxifene increased myelin thickness of remyelinated axons to pre-lesion levels. Taken together, our findings indicate that prolonging the ISR protects remyelinating oligodendrocytes and promotes remyelination in the presence of inflammation, suggesting that ISR enhancement may provide reparative benefit to MS patients.

## INTRODUCTION

Multiple sclerosis (MS) is an autoimmune inflammatory disorder characterized by focal demyelinated lesions in the central nervous system (CNS) (Frohman et al. 2006; Yadav et al. 2015; Reich et al. 2018). Although current immunomodulatory therapies can reduce the frequency and severity of relapses, they have demonstrated limited impact on the progression of disease (Dargahi et al. 2017; Hauser and Cree 2020). Complementary strategies are therefore urgently needed to protect oligodendrocytes and promote repair of the CNS in order to slow or even stop the progression of MS (Hart and Bainbridge 2016; Rodgers et al. 2013; Way and Popko 2016).

Remyelination is the process of restoring demyelinated nerve fibers with new myelin, which involves the generation of new mature oligodendrocytes from oligodendrocyte precursor cells (OPCs) (Franklin and Ffrench-Constant 2008). Failure of myelin repair during relapsing-remitting MS leads to chronically demyelinated axons, which is thought to contribute to axonal degeneration and disease progression. (Franklin and Kotter 2008; Fancy et al. 2010; Chari 2007). The inflammatory environment in MS lesions is considered a major contributor to impaired remyelination (Starost et al. 2020). Strategies to enhance myelin regeneration could preserve axonal integrity and increase clinical function of patients. A number of small molecules have recently been described that promote OPC proliferation and/or enhance remyelination (Deshmukh et al. 2013; Mei et al. 2014; Rankin et al. 2019; Najm et al. 2015). Given that remyelination in MS occurs in lesions with ongoing inflammation, a remyelination model with an inflammatory environment is needed to determine the potential of remyelinating-promoting compounds for MS treatment.

The integrated stress response (ISR) plays a key role in the response of oligodendrocytes to CNS inflammation (Chen et al. 2019; Lin et al. 2005; Way et al. 2015). The ISR is a cytoprotective pathway that is activated by various cytotoxic insults, including inflammation (Costa-Mattioli and Walter 2020). The ISR is initiated by phosphorylation of eukaryotic translation initiation factor 2 alpha (p-eIF2α), by one of four known stress-sensing kinases, to diminish global protein translation and selectively allow for the synthesis of protective proteins. p-eIF2α is dephosphorylated by the protein phosphatase 1 (PP1)-DNA-damage inducible 34 (GADD34) complex to terminate the ISR upon stress relief (Zhang and Kaufman 2008). Sephin1, a recently-identified small molecule, disrupts PP1-GADD34 phosphatase activity and prolongs elevated p-eIF2α levels, thereby enhancing the ISR protective response (Das et al. 2015; Carrara et al. 2017). We previously showed that GADD34 deficiency or Sephin1 treatment protects oligodendrocytes, diminishes demyelination, and delays clinical symptoms in a mouse model of MS, experimental autoimmune encephalomyelitis (EAE) (Chen et al. 2019).

Here, we investigated the potential impact of genetic or pharmacological ISR enhancement on remyelinating oligodendrocytes and the remyelination process in the presence of inflammation. In these studies, we used the EAE model, as well as the cuprizone-induced demyelination model on *GFAP/tTA;TRE/IFN-γ* double-transgenic mice, which ectopically express interferon gamma (IFN-*γ*) in the CNS in a regulated manner (Lin et al. 2004; Lin et al. 2005). IFN-*γ* is a pleotropic T cell specific cytokine that stimulates key inflammatory aspects of MS (Lin et al. 2007; Lees and Cross 2007). Using these inflammatory demyelination models, we show that prolonging the ISR protects remyelinating oligodendrocytes and increases the level of remyelination. Moreover, we explored the effects of combining Sephin1 treatment with bazedoxifene (BZA) (Rankin et al. 2019) on remyelination. BZA has been shown to enhance OPC differentiation and CNS remyelination in response to focal demyelination (Rankin et al. 2019). We demonstrate that BZA increases the number of remyelinating oligodendrocytes and enhances remyelination in the presence of inflammation. When combined with Sephin1, BZA further facilitates the remyelinating process.

## RESULTS

### Sephin1 treatment enhances remyelination in late-stage EAE

To determine whether pharmacological prolongation of the ISR can enhance remyelination after inflammatory demyelination, we first examined C57BL/6J mice in the late stage of EAE. Sephin1 (8 mg/kg) or vehicle treatment was initiated in each EAE mouse on the day it reached the peak of disease (clinical score = 3) and was continued to the late stage of EAE, which resulted in diminished EAE disease severity in the final week of Sephin1 treatment (Figure 1A). Spinal cord axons were examined under electron microscope (EM). We sorted remyelinated axons by examining g-ratios (axon diameter / total fiber diameter) in the lumbar spinal cord white matter of EAE mice after vehicle or Sephin1 treatment. The presence of axons with thinner myelin sheaths and a higher g*-*ratio is considered the hallmark of remyelination (Duncan et al. 2017). A recent study indicated that axons in the spinal cord with a g-ratio greater than 0.8 are likely remyelinated axons in the EAE model (Mei et al. 2016). Our data demonstrated that the density of remyelinated axons (g > 0.8) was significantly higher in EAE mice treated with Sephin1 than those treated with vehicle (Figure 1 B, C), although no difference was detected in the total number of myelinated axons (Figure 1D). The data suggest that Sephin1 promotes remyelination in the neuroinflammatory environment of EAE.

**Figure 1.**
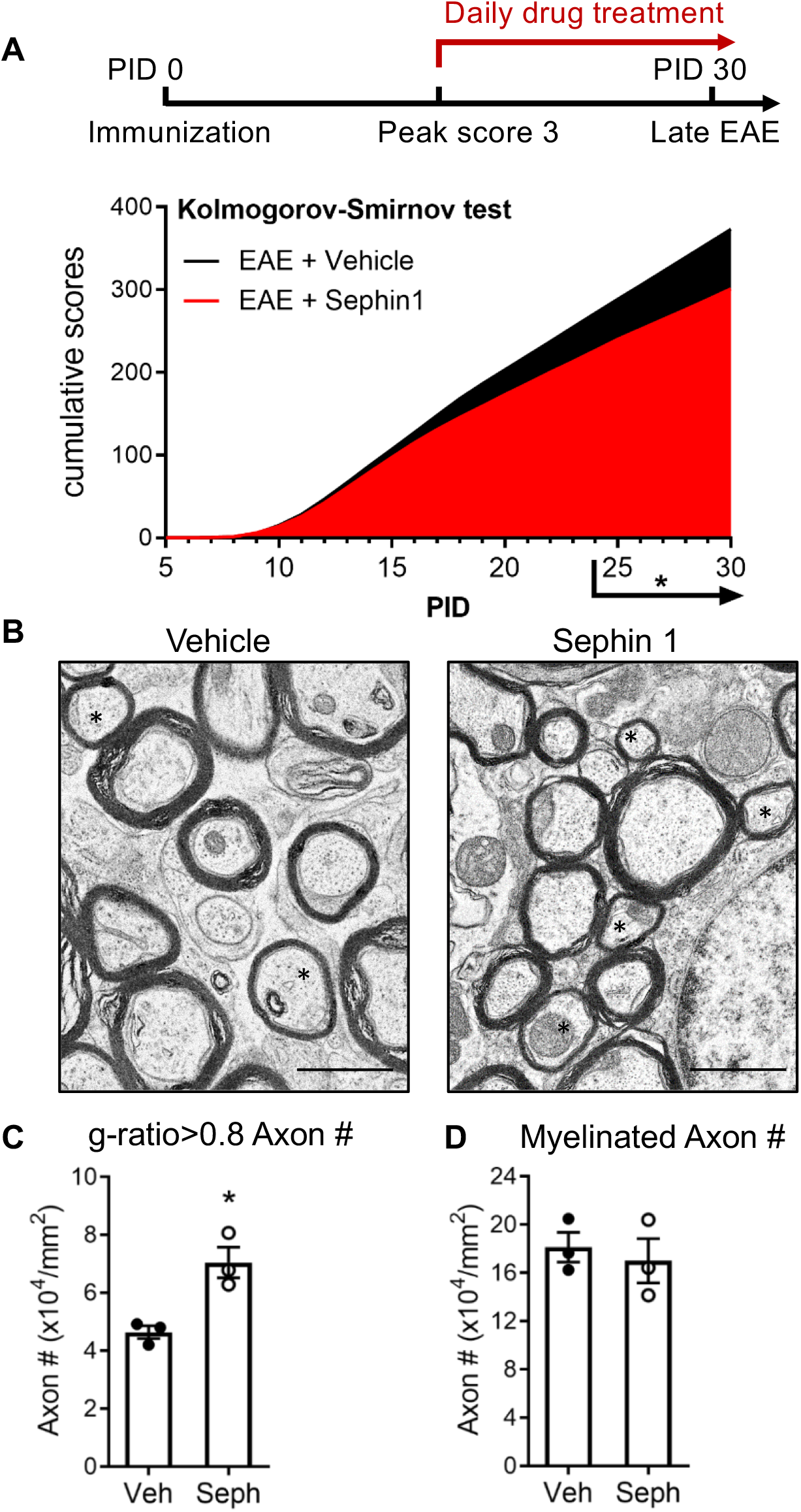
Sephin1 treatment enhances remyelination after inflammatory demyelination. (A) Cumulative clinical scores of C57BL/6J female mice immunized with MOG35-55/CFA to induce chronic EAE, treated with vehicle (*n =* 7) and 8 mg/kg Sephin1 (*n =* 7) from the peak disease. \**P* < 0.05. Significance based on Kolmogorov-Smirnov test. (B) Representative EM images of axons in the spinal cord white matter tracts of EAE mice treated with vehicle or Sephin1, starting at peak EAE clinical score. Remyelinated axons in the EAE spinal cord were identified by thinner myelin sheaths (*). Scale bar=1μm. (C) Density of myelinated axons with g-ratio <0.8 in the EAE spinal cord. (D) Density of myelinated axons in the EAE spinal cord. Data are presented as the mean ± SEM (n=3 mice/group). Over 300 axons were analyzed per mouse. \**P <* 0.05. Significance based on unpaired *t*-test.

### GADD34 deficiency protects remyelinating oligodendrocytes and enhances remyelination in the presence of IFN-γ

We have previously shown that in response to IFN-γ, a *GADD34* null mutation in myelinating oligodendrocytes increases levels of p-eIF2α, indicating a prolongation of the ISR, and promotes oligodendrocyte survival (Lin et al. 2008). We hence mated *GADD34* KO*;GFAP/tTA* mice with *GADD34* KO*;TRE/IFN-γ* mice to generate *GFAP/tTA;TRE/IFN-γ* double-transgenic mice homozygous for the *GADD34* mutation to prolong the ISR (Lin et al. 2008). *GFAP/tTA;TRE/IFN-γ* double-transgenic mice allow for ectopic release of IFN-γ to the CNS in a doxycycline-dependent manner (Lin et al. 2004; Lin et al. 2006; Lin et al. 2008). Expression of the tetracycline-controlled transactivator (tTA) is driven by the astrocyte-specific transcriptional regulatory region of the *GFAP* gene. In the *TRE/IFN-γ* mice, the IFN-γ cDNA is transcriptionally controlled by the tetracycline response element (TRE) (Figure 2A). The expression of the IFN-γ transgene is repressed in the *GFAP/tTA;TRE/IFN-γ* mice by providing water treated with doxycycline (Dox) from conception. At six weeks of age, transgenic mice were taken off Dox water to induce CNS expression of IFN-γ (IFN-γ+) and placed on a diet of 0.2% cuprizone chow (Lin et al. 2006; Lin et al. 2004). After five weeks of cuprizone exposure, mice were placed back on a normal diet for up to three weeks to allow remyelination to occur. Control mice received Dox water throughout the study to repress CNS expression of IFN-γ (IFN-γ-) (Figure 2B). Mice with the highest level of IFN-γ expression in the CNS were selected by isolating the cerebellum of each mouse and using real-time reverse transcription (RT)-PCR to determine IFN-γ expression levels. We found that at five weeks of cuprizone exposure (W5) and during remyelination (W8), removal of Dox significantly increased the levels of IFN-γ in the cerebellum of mice (Figure 2 – figure supplement 1A).

**Figure 2.**
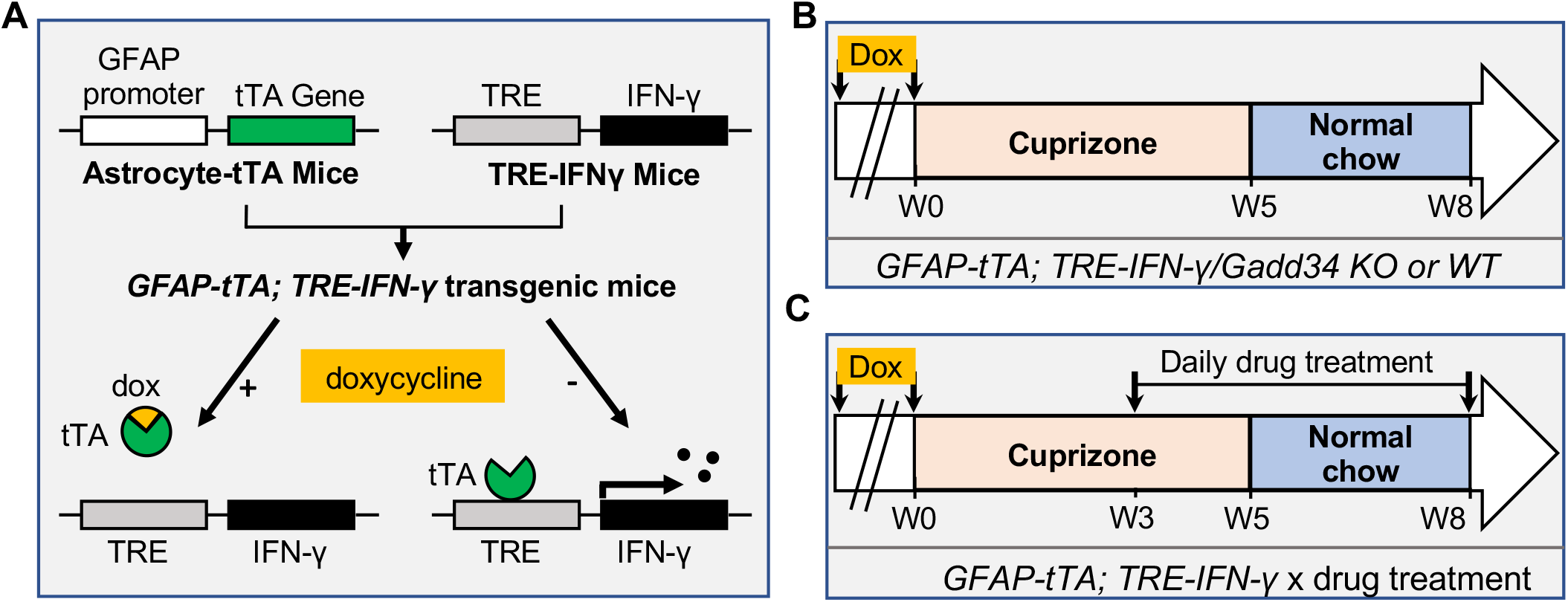
Schematics of *GFAP/tTA;TRE/IFN-γ* double-transgenic mouse model of cuprizone demyelination/remyelination. (A) *GFAP/tTA* mice are mated with *TRE/IFN-γ* mice to produce double-positive animals. When these mice are maintained on doxycycline, the expression of the IFN-γ is repressed. When they are released from doxycycline, IFN-γ is expressed in the CNS. (B) Cuprizone demyelination/remyelination model of *GFAP/tTA*;*TRE/IFN-γ*/*GADD34 KO* or *WT*. Doxycycline (Dox) is removed and cuprizone chow is added when mice are at six-week-old (W0). After 5 weeks of cuprizone exposure (W5), mice were placed back on normal chow for up to three weeks to allow remyelination. (C) Cuprizone demyelination/remyelination model of *GFAP/tTA*;*TRE/IFN-γ* with designed treatment. Drug treatment is started at 3 weeks of curpzione exposure (W3) and lasts to the end of remyelination (W8). The following figure supplement is available for figure 2. **Figure supplement 1.** *GFAP/tTA;TRE/IFN-γ* mice express IFN-γ after release from doxycycline.

We next investigated whether the *GADD34* mutation promotes oligodendrocyte survival in the presence of IFN-γ during demyelination/remyelination in the cuprizone model. It has been demonstrated that cuprizone-fed mice exhibit apoptotic death of oligodendrocytes and demyelination in the corpus callosum. Complete remyelination spontaneously occurs a few weeks after the cuprizone challenge is terminated (Matsushima and Morell 2001). We found no significant difference in the number of cells labeled with the mature oligodendrocyte marker ASPA between *GADD34* wildtype (WT) and *GADD34* KO of *GFAP/tTA;TRE/IFN-γ* double-transgenic mice before cuprizone exposure (Figure 3A, B). After five weeks on cuprizone chow, both *GADD34* WT and KO double-transgenic mice presented similarly with a significantly reduced number of ASPA+ mature oligodendrocytes in the presence of IFN-γ (IFN-γ+) (WT, *p<0.001*; KO, *p<0.05*) (Figure 3A, B). During the remyelination period, ASPA+ oligodendrocytes reappeared and reached approximately 800/mm^2^ in both IFN-γ-repressed *GADD34* WT and KO mice (IFN-γ-) after three weeks of normal chow. In the presence of IFN-γ (IFN-γ+), this number dropped in the lesions of WT double-transgenic mice to roughly 500/mm^2^ of ASPA+ cells (*p<0.05*) (Figure 3A, C), which is consistent with our previous finding that induction of IFN-γ expression in the CNS suppresses the repopulation of oligodendrocytes following cuprizone-induced oligodendrocyte toxicity (Lin et al. 2006). Compared to *GADD34* WT, *GFAP/tTA;TRE/IFN-γ* mice with the *GADD34* mutation had significantly increased numbers of ASPA+ oligodendrocytes during remyelination in the presence of IFN*-γ* (IFN-γ+/W8) (*p<0.01*) (Figure 3A, C). Myelin status in these mice was also evaluated with EM. After five weeks of cuprizone chow, a significant number of axons exhibited myelin sheath loss in control mice (IFN-γ*-*), compared to 0 weeks (WT, *p<0.0001*; KO, *p<0.001*) (Figure 4A, C). Similarly, both WT and KO double-transgenic mice presented markedly reduced myelinated axons in the presence of IFN*-*γ (IFN-γ+) (WT: 25.2±3.0%, *p<0.0001*; KO: 24.4±2.7%, *p<0.0001*). Nevertheless, no difference in the percentage of myelinated axons was observed between IFN-γ+ and IFN-γ-groups (Figure 4A, C). At three weeks after cuprizone withdrawal, both IFN-γ-repressed WT and KO mice (IFN-γ-) showed spontaneous remyelination (WT: 42.7±6.2%; KO: 42.6±4.7%) (Figure 4B, D). Meanwhile, remyelination was significantly suppressed in the corpus callosum of IFN*-*γ-expressing WT mice (WT: 29.4±2.9%) (*p<0.05*), but not in that of IFN*-*γ-expressing KO mice (KO: 49.1±8.8%) (Figure 4B, D). Together, these data suggest that prolonging the ISR has no impact in the absence of IFN-γ mice, but it results in increased repopulation of oligodendrocytes and remyelination following demyelination in the presence of IFN-γ.

**Figure 3.**
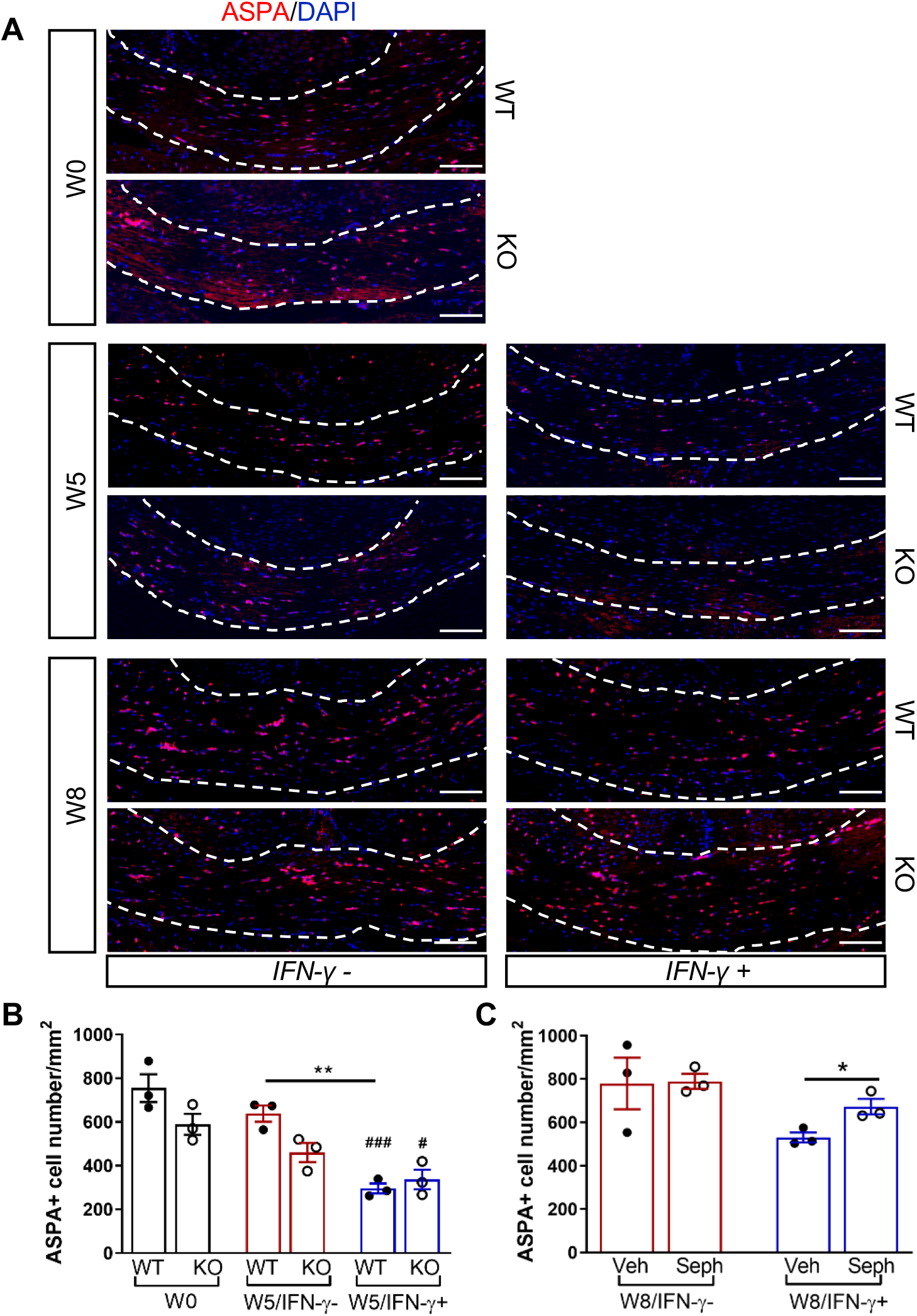
GADD34 deficiency protects remyelinating oligodendrocytes in the presence of IFN-γ. The corpora callosa of *GFAP/tTA*;*TRE/IFN-γ*/*GADD34 KO* or *WT* were taken at W0, W5 and W8. (A) Immunofluorescent staining for ASPA (a mature oligodendrocyte marker) and DAPI (nuclei). Scale bar=100μm. (B) Quantification of cells positive for ASPA in the corpus callosum areas at W0 and W5 in the absence (IFN-γ-) or presence of IFN-γ (IFN-γ+). Data are presented as the mean ± SEM (n=3 mice/group). \*\**P <* 0.01, ^#^*P <* 0.05 (#vs W0/WT), ^###^*P <* 0.001 (# vs W0/KO). Significance based on ANOVA. (C) Quantification of cells positive for ASPA in the corpus callosum areas at W8 in the absence or presence of IFN-γ. Data are presented as the mean ± SEM (n=3 mice/group). \**P <* 0.05, \*\**P <* 0.01. Significance based on ANOVA.

**Figure 4.**
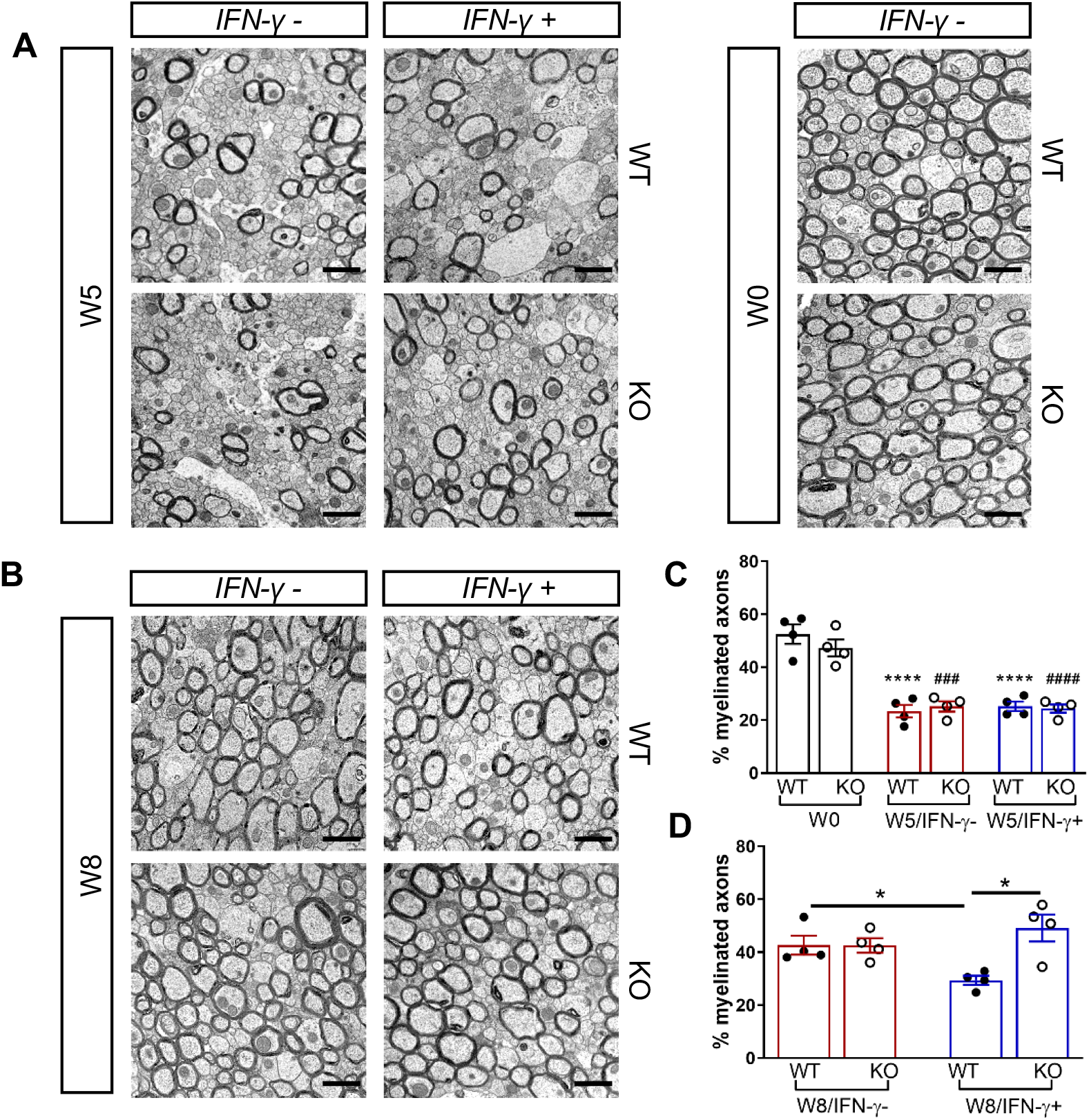
GADD34 deficiency enhances remyelination in the presence of IFN-γ. The corpora callosa of *GFAP/tTA*;*TRE/IFN-γ*/*GADD34 KO* or *WT* were harvested for EM processing. (A) Representative EM images of axons in the corpus callosum at W0 and W5. Scale bar=1μm. (B) Representative EM images of axons in the corpus callosum at W8. Scale bar=1μm. (C) Percentage of remyelinated axons at W0 and W5. (D) Percentage of remyelinated axons at W8. Data are presented as the mean ± SEM (n=4 mice/group). \**P <* 0.05, \*\**P <* 0.01, ****P<0.0001. Significance based on ANOVA.

### Sephin1 treatment protects remyelinating oligodendrocytes and enhances remyelination in the presence of IFN-γ

We have previously demonstrated that Sephin1 treatment can prolong the ISR in primary oligodendrocytes exposed to IFN-γ, shown by prolonged elevated eIF2α phosphorylation levels, (Chen et al. 2019). In addition, Sephin1 protects mature oligodendrocytes against inflammation in the EAE model (Chen et al. 2019). Therefore, using Sephin1, we tested the ability of pharmacological enhancement of the ISR to protect remyelinating oligodendrocytes from inflammatory insults in the cuprizone model. Similarly to the GADD34 experiment, six-week-old *GFAP/tTA;TRE/IFN-γ* double transgenic mice that had been on Dox water since conception were either kept on (IFN-γ-) or removed from Dox (IFN-γ+) and fed with a chow containing 0.2% cuprizone for five weeks. Mice were then placed back on a normal diet for three weeks. Sephin1 (8 mg/kg) or vehicle was administered daily to the mice by intraperitoneal injections beginning three weeks after the start of the cuprizone diet. This time point represents the peak of oligodendrocyte loss during cuprizone-mediated demyelination (Matsushima and Morell 2001) (Figure 2C).

We first examined IFN-γ expression in the double-transgenic mice, which showed that mice removed from Dox (IFN-γ+) expressed significantly higher levels of IFN-γ in the cerebellum than mice on Dox water (IFN-γ-) at five weeks and eight weeks of cuprizone treatment in all groups (Figure 2 – figure supplement 1B). Next, we investigated the number of mature oligodendrocytes in the corpus callosum by immunofluorescent staining of ASPA. Three weeks of cuprizone exposure led to a significant decrease of the ASPA+ oligodendrocytes in both IFN-γ-(*p<0.01*) and IFN-γ+ (*p<0.01*) groups (Figure 5A, B). After five weeks of cuprizone chow, we observed a further reduction in the number of oligodendrocytes in the presence of IFN*-*γ (*p*<0.01), although we found no difference between vehicle and Sephin1 treatment (veh: 315±44/mm^2^; Seph 394±67/mm^2^) (Figure 5A, B). At three weeks after cuprizone withdrawal, the density of oligodendrocytes returned to the initial levels seen at week 0 (W0) in both vehicle- and Sephin1-treated mice in the absence of IFN-γ. Activation of IFN*-γ* expression by removing Dox (IFN-γ+) suppressed the repopulation of ASPA+ oligodendrocytes in the vehicle treated mice, but Sephin1 treatment significantly increased the survival of these cells (veh: 531±32/mm^2^, Seph: 672±52/mm^2^, *p<0.05*) (Figure 5A, C). In addition, we observed approximately 50% reduction in the percentage of myelinated axons after five weeks of cuprizone exposure in both IFN-γ- and IFN-γ+ double-transgenic mice *(p<0.0001)* (Figure 6A, C). Similar to the changes in oligodendrocyte numbers, our data demonstrated that during remyelination, CNS delivery of IFN*-γ* resulted in diminished numbers of remyelinated axons (*p<0.05*), while mice treated with Sephin1 exhibited a significantly higher percentage of remyelinated axons (veh: 30.3±3.3%, Seph: 43.2±4.6%, *p<0.01*) (Figure 6B, D). These data indicate that although the pharmacological enhancement of the ISR with Sephin1 has no impact in the absence of IFN-γ, it can provide protection to remyelinating oligodendrocytes and promote remyelination in the presence of IFN-γ.

**Figure 5.**
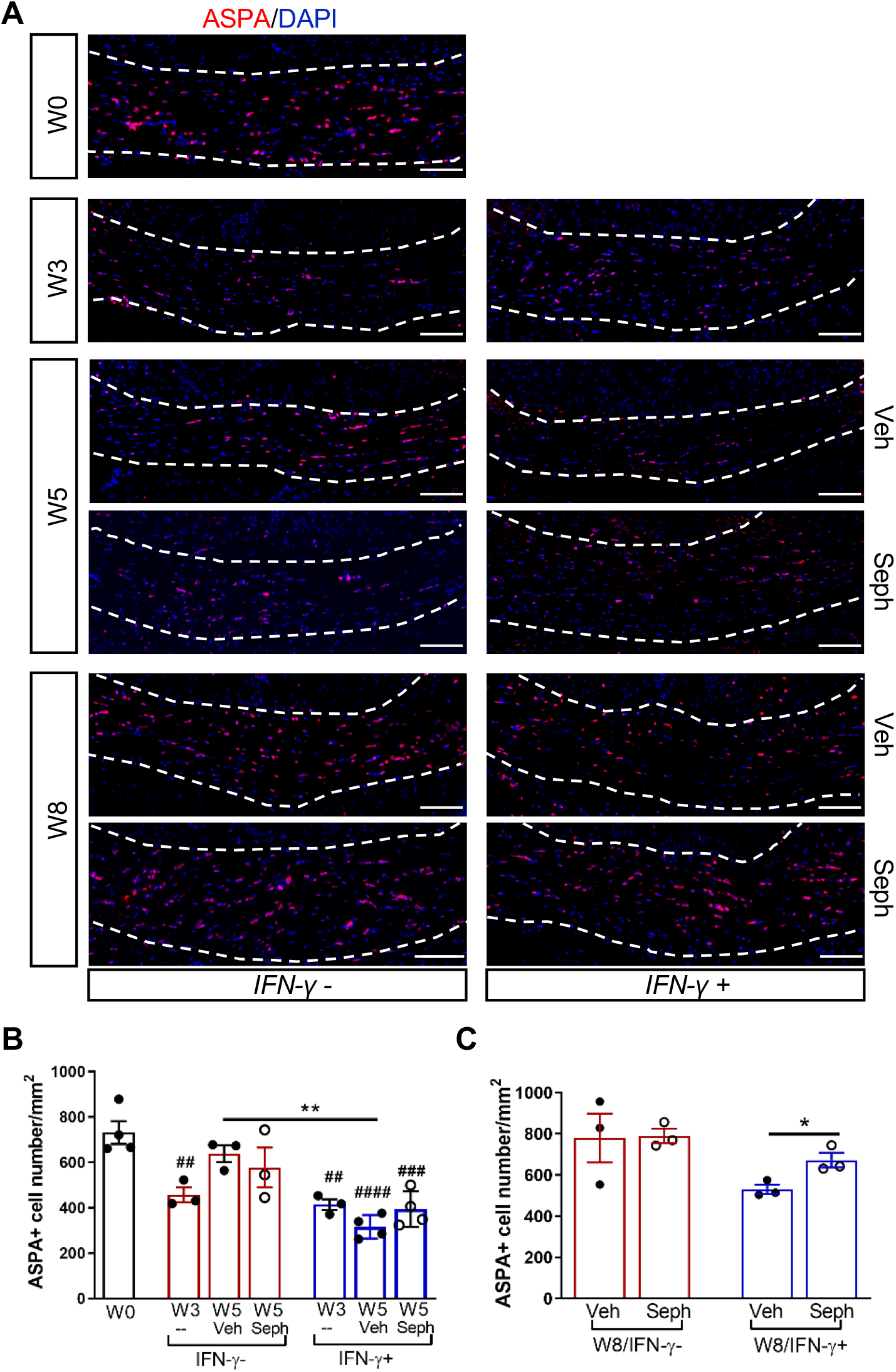
Sephin1 protects remyelinating oligodendrocytes in the presence of IFN-γ. The corpora callosa of *GFAP/tTA*;*TRE/IFN-γ* were taken at W0 and W3 prior to any treatment as well as after either vehicle or Sephin1 treatment at W5 and W8. (A) Immunofluorescent staining for ASPA (a mature oligodendrocyte marker) and DAPI (nuclei). Scale bar=100μm. (B) Quantification of cells positive for ASPA in the corpus callosum areas at W0, W3 and W5 in the absence (IFN-γ-) or presence of IFN-γ (IFN-γ+). Data are presented as the mean ± SEM (n=3-4 mice/group). \*\**P <* 0.01, ^##^*P <* 0.01, ^###^*P <* 0.001, ^####^*P <* 0.0001 (#vs W0). Significance based on ANOVA. (C) Quantification of cells positive for ASPA in the corpus callosum areas at W8 in the absence or presence of IFN-γ. Data are presented as the mean ± SEM (n=3 mice/group). \**P <* 0.05. Significance based on ANOVA.

**Figure 6.**
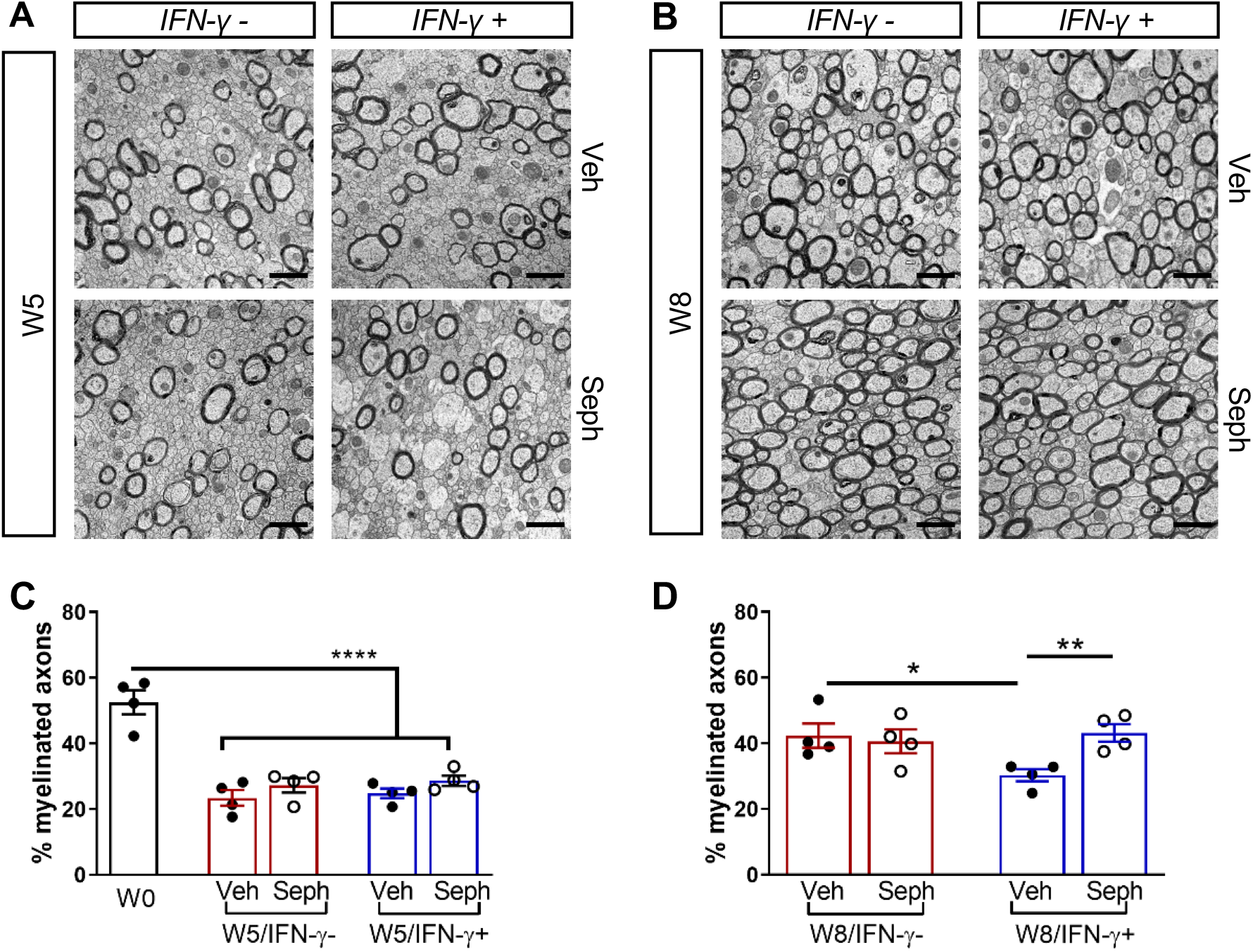
Sephin1 enhances remyelination in the presence of IFN-γ. The corpora callosa of *GFAP/tTA*;*TRE/IFN-γ*/*GADD34 KO* or *WT* were harvested for EM processing. (A) Representative EM images of axons in the corpus callosum at W5. Scale bar=1μm. (B) Representative EM images of axons in the corpus callosum at W8. Scale bar=1μm. (C) Percentage of remyelinated axons at W0 and W5. (D) Percentage of remyelinated axons at W8. Data are presented as the mean ± SEM (n=4 mice/group). \**P* < 0.05, \*\**P <* 0.01, \*\*\*\**P*<0.0001. Significance based on ANOVA.

### GADD34 deficiency or Sephin1 treatment does not affect OPC proliferation or microglial recruitment following cuprizone demyelination/remyelination

Recruitment of OPCs is required for spontaneous remyelination, during which OPCs proliferate and migrate to the demyelinated area to mature into functional oligodendrocytes (Huang and Franklin 2011; Franklin and Ffrench-Constant 2008). Using the OPC marker PDGFRα and the proliferation marker Ki67, we noted the appearance of PDGFRα+ OPCs and proliferating OPCs (PDGFRα+/Ki67+) near the demyelinated lesions (W5) and remyelinated areas (W8) in the corpus callosum of double-transgenic mice, which was not affected by GADD34 deficiency (Figure 7 A–C) or Sephin1 treatment (Figure 7 D–F). Interestingly, at week 5 of cuprizone exposure, the induction of IFN-*γ* by Dox removal (IFN-γ+) diminished the number of proliferating OPCs (PDGFRα+/Ki67+) in vehicle-treated mice (*p<0.05*), but no difference between vehicle and Sephin1 was found in the number of OPCs in the presence of IFN-γ (Figure 7 D, E).

**Figure 7.**
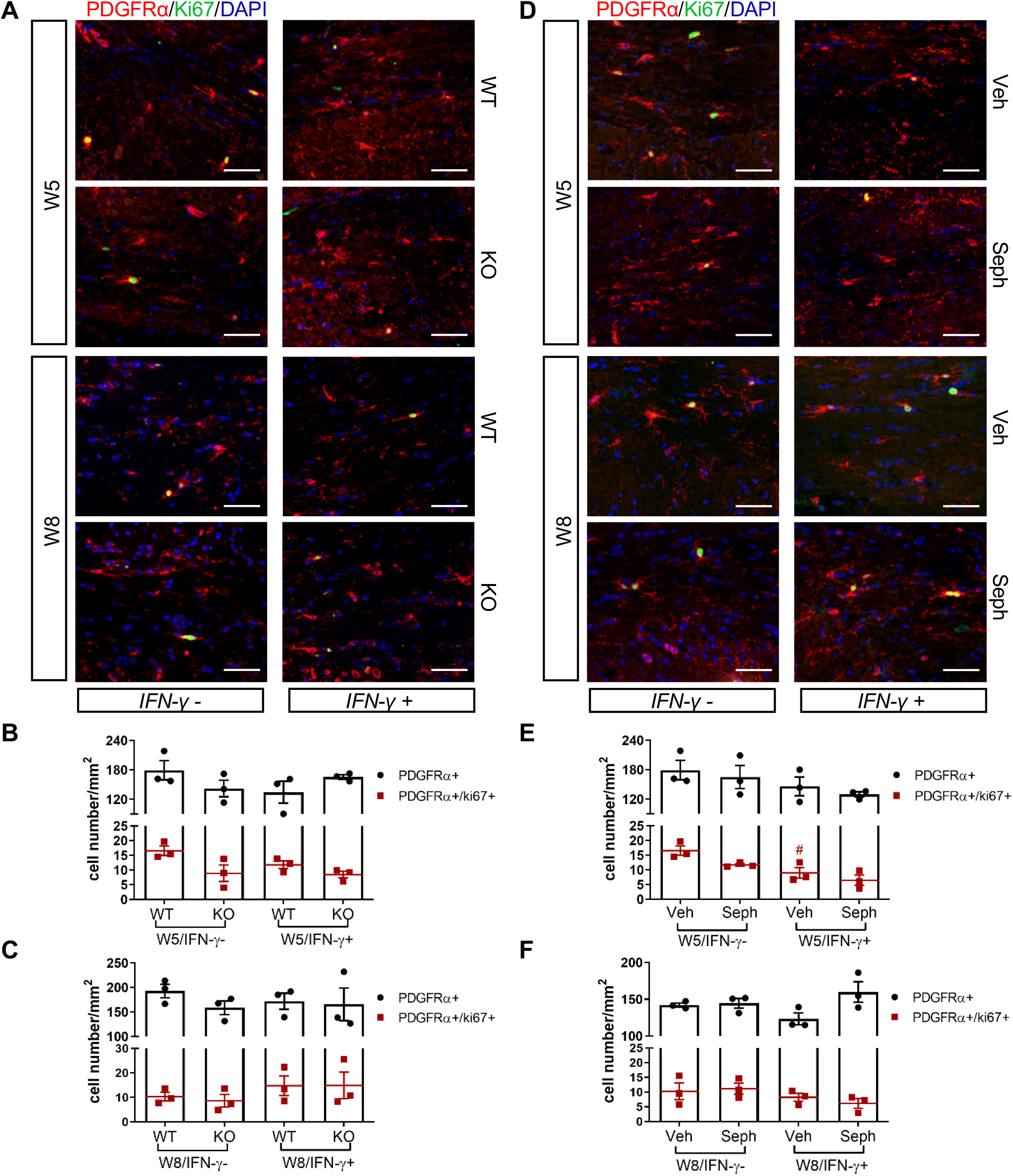
GADD34 deficiency or Sephin1 does not affect OPC proliferation during remyelination. (A) Immunofluorescent staining for PDGFRα (an OPC marker), Ki67 and DAPI (nuclei) from the corpus callosum of *GFAP/tTA*;*TRE/IFN-γ*/*GADD34 KO* or *WT* was taken at W5 and W8. Scale bar=50μm. Quantification of cells positive for PDGFRα and cells positive for both PDGFRα and Ki67 in the corpus callosum areas of *GFAP/tTA*;*TRE/IFN-γ*/*GADD34 KO* or *WT* in the absence (IFN-γ-) or presence of IFN-γ (IFN-γ+) at W5 (B) and W8 (C). Data are presented as the mean ± SEM (n=3 mice/group). (D) Immunofluorescent staining for PDGFRα, Ki67 and DAPI from the corpus callosum of *GFAP/tTA*;*TRE/IFN-γ* was taken after either vehicle or Sephin1 treatment at W5 and W8. Quantification of cells positive for PDGFRα and cells positive for both PDGFRα and Ki67 in the corpus callosum areas of *GFAP/tTA*;*TRE/IFN-γ* treated with treatment in the absence (IFN-γ-) or presence of IFN-γ (IFN-γ+) at W5 (E) and W8 (F). Data are presented as the mean ± SEM (n=3 mice/group). #*P <* 0.05 (vs. veh from W5/IFN-γ-). Significance based on ANOVA. The following figure supplements are available for figure 7. **Figure supplement 2.** GADD34 deficiency does not affect microglial activation during remyelination in the presence of IFN-γ. **Figure supplement 3.** Sephin1 treatment does not affect microglial activation during remyelination in the presence of IFN-γ.

Studies have indicated that recruitment of microglia is required for myelin clearance and efficient initiation of remyelination (Neumann et al. 2009; Voss et al. 2012; Gudi et al. 2014). To examine the activation of microglia, we immuno-stained sections of corpus callosum with the microglia marker IBA1 and quantified the density of these cells in the lesions. A significantly stronger microgliosis was observed in the corpus callosum of double-transgenic mice after five weeks of cuprizone chow (W5) and activated microglia persisted after three weeks of remyelination (W8) (Figure 7 – figure supplement 2 and Figure 7 – figure supplement 3). Interestingly, in the group of GADD34 KO and of Sephin1 treatment, the number of IBA1+ cells at W8 was significantly increased in the presence of IFN-γ than those in the absence of IFN-γ (Figure 7 – figure supplement 2 A, C and Figure 7 – figure supplement 3 A, C), but we did not detect a statistical change in the density of IBA1+ cells between WT mice and KO or between vehicle and Sephin1 treatment (Figure 7 – figure supplement 2 A, C and Figure 7 – figure supplement 3 A, C).

### Combined treatment of Sephin1 and BZA enhanced remyelination in the presence of IFN-γ

BZA is a selective estrogen receptor modulator (SERM) that is currently FDA-approved to be used in combination with conjugated estrogen in menopausal women (McKeand et al. 2014). Recently, Rankin et al. showed that BZA significantly enhances OPC differentiation and accelerates remyelination in the lysolecithin focal demyelination model (Rankin et al. 2019). Given that the enhancement of the ISR with Sephin1 provided protection to remyelinating oligodendrocytes in the presence of inflammation (Figure 5), we next assessed whether the combined treatment of Sephin1 and BZA is more effective in enhancing remyelination than either treatment alone (Figure 2C). We found that Sephin1, BZA, and combined Sephin1/BZA treatment significantly increased ASPA+ oligodendrocyte numbers in the corpus callosum during remyelination in the presence of IFN-γ (W8/ IFN-γ+) (Figure 8 A, C). Accordingly, compared with vehicle controls, a significant increase in the percentage of myelinated axons was noted in the groups treated with Sephin1, BZA, and combined Sephin1/BZA treatment (Figure 8 B, D). Nonetheless, no difference between treatment groups was observed either for oligodendrocyte survival or number of remyelinated axons, although the number of remyelinated axons in treatment groups reached pre-lesion levels (Figure 8 B, D). In addition to measuring the, percentage of myelinated axons, we also examined myelin thickness using EM analysis of gratios after remyelination. The vehicle-treated group (W8/IFN-γ+) demonstrated a significantly higher g-ratio (0.815±0.012) than those at W0 (IFN-γ-). In contrast, corpora callosa axons in mice receiving the combination treatment of Sephin1 and BZA (g-ratio: 0.778±0.006) reached myelin thickness levels comparable with pre-lesion axons (Figure 8 B, E).These data suggest that the combination treatment restored myelin thickness to baseline levels. Moreover, g-ratios of myelinated axons were significantly lower in the combination treatment than the vehicle-treated group (*p*<0.01) or each of the single treatment groups (*p*<0.05 vs Sephin1, *p*<0.05 vs BZA), indicating that the combined therapy enhanced the remyelination response.

**Figure 8.**
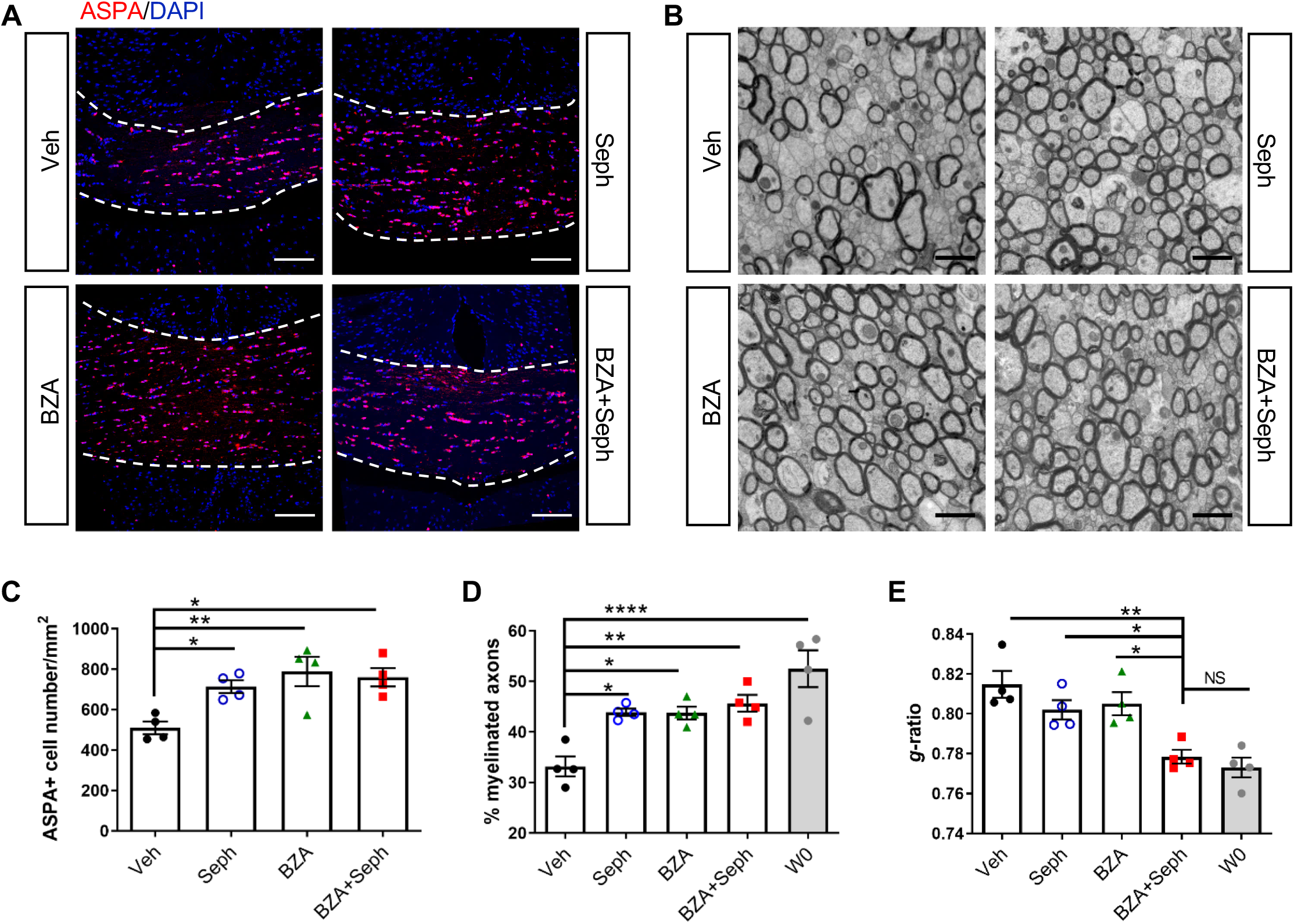
Combined treatment of Sephin1 and BZA further enhance remyelination. *GFAP/tTA*;*TRE/IFN-γ* mice were released from Dox at W0 and given cuprizone chow for 5 weeks. Treatment with vehicle, Sephin1, BZA or combined BZA and Sephin1 was started at W3, and the corpus callosum was harvested at W0 and after 3 weeks of remyelination (W8). (A) Immunofluorescent staining for ASPA and DAPI. Scale bar=100μm. (B) Quantification of cells positive for ASPA in the corpus callosum areas at W0 and W8 in the presence of IFN-γ. (C) Representative EM images of axons in the corpus callosum at W8 in the presence of IFN-γ. Scale bar=1μm. Data are presented as the mean ± SEM (n=4 mice/group). \**P <* 0.05, ^**^*P <* 0.01, ^****^*P*<0.0001. Significance based on ANOVA.

## DISCUSSION

Endogenous remyelination plays a crucial role in preserving axons and restoring neuronal function, but is frequently insufficient in MS. The adverse extracellular environment of MS lesions is thought to contribute to remyelination failure (Chew et al. 2005; Frischer et al. 2015; Kuhlmann et al. 2008; Franklin and Ffrench-Constant 2008). Efforts to re-populate oligodendrocytes that remyelinate neuronal axons within the CNS will advance reparative therapeutics for MS (Franklin and Kotter 2008; Fancy et al. 2010; Chari 2007). Although many agents identified from high-throughput screening studies appear to accelerate remyelination, it is unclear whether they are capable of initiating repair and restoring neuronal function in a complex inflammatory environment (Deshmukh et al. 2013; Mei et al. 2016). A recent study, which investigated the differentiation of induced pluripotent stem cell-derived oligodendrocytes from MS patients, indicated that the inflammation in MS lesions is a major contributor to impaired remyelination, and many remyelinating agents failed to restore impaired differentiation caused by inflammation (Starost et al. 2020). Our mouse model used here, which combines primary demyelination induced by cuprizone with CNS delivery of IFN-γ, provides a unique platform to facilitate the assessment of remyelination therapies in the setting of inflammation. In our previous studies, we demonstrated that prolonging the ISR provides protection to mature oligodendrocytes, which maintain the myelin sheath, from inflammatory attack (Way and Popko 2016; Way et al. 2015; Lin et al. 2008). Herein we investigated whether a similar enhancement of the ISR would provide protection to remyelinating oligodendrocytes and promote the repair of demyelinated axons in a neuroinflammatory environment.

We have previously shown that a prolonged ISR response protects mature oligodendrocytes during the peak of disease in EAE in a manner not related to direct immunomodulation (Chen et al. 2019). Here, we demonstrated that Sephin1 treatment, when initiated at the peak of disease, facilitated the clinical recovery of the EAE animals, which correlated with enhanced remyelination. These results motivated us to examine the remyelination enhancing potential of an augmented ISR in a more controlled model of CNS remyelination.

To further investigate the potential of the ISR to confer protection to remyelinating oligodendrocytes and remyelination during inflammation, we utilized an inducible double-transgenic mouse model (*GFAP/tTA;TRE/IFN-γ*) to temporally deliver IFN-γ to the CNS in the cuprizone model (Lin et al. 2006). The T cell cytokine IFN-γ is a critical inflammatory mediator of MS/EAE pathogenesis (Lees and Cross 2007; Ottum et al. 2015). In addition, it has been suggested that IFN-γ could induce immune transitions of OPCs and oligodendrocytes to an inflammatory state with major histocompatibility complex (MHC)-I and MHC-II presentation (Kirby and Castelo-Branco 2020). IFN-γ delivery is required as cuprizone-induced demyelination/remyelination occurs in the absence of an adaptive immune response and thus does not reflect the inflammatory environment in the MS lesions. Importantly, the level of IFN-γ present in the CNS of *GFAP/tTA;TRE/IFN-γ* double transgenic mice is approximately equivalent to that of EAE mice (data not shown), which suggests that Dox-controlled IFN-γ release is within the range observed in inflammatory demyelination. Our previous studies with *GFAP/tTA;TRE/IFN-γ* double-transgenic mice demonstrated that ectopic expression of IFN-γ in the CNS causes a reduction in the number of myelinating oligodendrocytes and subsequent hypomyelination during development, and results in diminished remyelination following cuprizone-induced oligodendrocyte toxicity (Lin et al. 2008; Lin et al. 2006). In our current study we found that CNS expression of IFN-γ suppresses the repopulation of oligodendrocytes and results in diminished numbers of remyelinated axons following cuprizone-induced demyelination. When oligodendrocytes are exposed to IFN-γ, the ISR is activated, which could play a key role in protecting remyelinating oligodendrocytes against the inflammatory environment (Lin et al. 2007; Lin et al. 2008; Chen et al. 2019; Lin et al. 2005). Indeed, GADD34 deficiency or Sephin1 treatment, both of which prolong the ISR, significantly increased the number of remyelinating oligodendrocytes and myelinated axons in the presence of IFN-γ. Nevertheless, the enhancement of the ISR had no effect on remyelination in the absence of IFN-γ. This is consistent with a recent study that showed that guanabenz, a drug closely related to Sephin1 (Das et al. 2015), did not improve remyelination in the non-inflammatory cuprizone model of demyelination (Thompson and Tsirka 2020).

The genetic and pharmacologic approaches used here to modulate the ISR do not target specific cell types in the CNS. The increased numbers of remyelinating oligodendrocytes that result from an enhanced ISR might originate from increased OPC survival in the lesion or from more efficient OPC differentiation to mature oligodendrocytes. We observed that proliferating OPC numbers at the peak of cuprizone-induced demyelination were diminished by the presence of IFN-γ, which is consistent with a previous finding that showed that IFN-γ predisposed OPCs to apoptosis (Chew et al. 2005). Nevertheless, prolonging the ISR by the *GADD34* mutation or by Sephin1 treatment did not alter the number of proliferating OPCs in the presence of inflammation. Therefore, it is likely that the benefits of ISR enhancement on remyelination in the presence of inflammation are primarily due to the protection of actively myelinating oligodendrocytes, similar to what we have observed during developmental myelination (Lin et al. 2005).

The FDA-approved SERM, BZA, has been shown to be capable of enhancing OPC differentiation and remyelination independently of its estrogen receptor (Rankin et al. 2019). We show here that BZA can promote remyelination in the presence of the inflammatory cytokine IFN-γ, which provides valuable preclinical support to the current MS clinical trial of BZA as a remyelination-enhancing agent (ClinicalTrials.gov Identifier: NCT04002934). Encouragingly, we found that the combination treatment of BZA and Sephin1 resulted in more axons with thicker myelin, indicating a more substantial recovery. The myelin sheaths of remyelinated axons are uniformly thin regardless of axon diameter, but the underlying mechanism that controls the thickness of remyelinated axons is unknown (Fancy et al. 2011). Although previous studies have shown that the transgenic overexpression of neuregulin (*Nrg*) or the conditional knockout of *PETN* increases myelin sheath thickness in developmental myelination, the remyelinated myelin sheaths remain thin in these models (Brinkmann et al. 2008; Harrington et al. 2010). Increased myelin thickness after the combination treatment of BZA and Sephin1 in our inflammatory de/remyelination model suggests that remyelination can be more efficiently induced by using therapies targeting OPC differentiation in combination with therapies that protect the remyelinating oligodendrocytes against the inflammatory environment.

Taken together, our current study demonstrates that the ISR modulator Sephin1 protects remyelinating oligodendrocytes in the presence of an inflammatory environment, leading to remarkably improved remyelination. In our previous EAE study, we showed that Sephin1 protects mature oligodendrocytes, myelin and axons, in addition to indirectly dampening the CNS inflammation that drives EAE (Chen et al. 2019). Combining current and previous findings, we believe that Sephin1 or similar ISR enhancing strategies will likely provide significant therapeutic benefit to MS patients.

## MATERIALS AND METHODS

### Animal study

Six-week-old female C57BL/6J mice were purchased from the Jackson Laboratory (Bar Harbor, ME, USA). The mice were allowed to acclimate for 14 days prior to the EAE experiment. The *GFAP/tTA* mice on the C57BL/6J background were mated with the *TRE/IFN-γ* mice on the C57BL/6J background to produce *GFAP/tTA; TRE/IFN-γ* double-transgenic mice (Lin et al. 2006; Lin and Popko 2009; Lin et al. 2004). Moreover, *GFAP/tTA* mice or *TRE/IFN-γ* were crossed with *GADD34* −/− mice, and the resulting progeny *GFAP/tTA*; *GADD34* −/− were crossed with *TRE/IFN-γ; GADD34* −/− to obtain double-transgenic mice that were homozygous for the *GADD34* mutation. To prevent transcriptional activation of IFN-γ, 0.05 mg/ml doxycycline (Dox, Sigma-Aldrich, St. Louis, MO, USA) was added to the drinking water and provided *ad libitum* from conception. Animals used in this study were housed under pathogen-free conditions at controlled temperatures and relative humidity with a 12/12-h light/dark cycle and free access to pelleted food and water. All animal experiments were conducted in accordance with ARRIVE guidelines and in complete compliance with Animal Care and Use Committee guidelines of the University of Chicago and Northwestern University.

The mice were randomly assigned to the different experimental groups. Sephin1 (free base) was purchased from Apexbio (Houston, TX, USA), and BZA (bazedoxifene acetate) from Sigma-Aldrich. Stock solutions of Sephin1 (24 mg/ml) and BZA (10mg/ml) in dimethyl sulfoxide (DMSO) were stored at −20C°. Final solutions were prepared in sterile 0.9% NaCl (DMSO concentration: 1%) for animal treatment.

### EAE immunization and treatment

EAE was induced in 7-week-old female C57BL/6J mice by subcutaneous flank administration of 200 μg MOG_35-55_ peptide emulsified with complete Freund’s adjuvant (CFA) (MOG_35-55_/CFA) (BD Biosciences, San Jose, CA, USA) *supplemented with inactive, Mycobacterium tuberculosis* H37Ra (BD Biosciences). Intraperitoneal (i.p.) injections of 200 ng pertussis toxin (List Biological Laboratories) were given immediately after administration of the MOG emulsion and again 48 hours later. Mice were blindly scored daily for clinical signs of EAE as follows: 0 = healthy, 1 = flaccid tail, 2 = ataxia and/or paresis, 3 = paralysis of hindlimbs and/or paresis of forelimbs, 4 = tetraparalysis, 5 = moribund or death. Mouse groups were randomized during the treatment. Mice were injected intraperitoneally with Sephin1 or the equivalent amount of vehicle (1% DMSO in 0.9% NaCl) daily starting from the peak of disease (score = 3). Lumbar spinal cords were collected at PID31.

### Cuprizone administration

To induce demyelination, *GFAP/tTA;TRE/IFN-γ and GFAP/tTA;TRE/IFN-γ;GADD34−/−* double transgenic mice were fed with a 0.2% cuprizone diet (Envigo, Madison, WI, USA) starting from six-weeks-old. Dox water was discontinued at the time of cuprizone treatment (Week 0, W0). Cuprizone feeding lasted five weeks and then mice were placed back on normal chow for up to three weeks to allow remyelination to occur. Control mice were maintained on Dox water during the entire experiment. 8 mg/kg of Sephin1 (i.p.) or 10 mg/kg of BZA (gavage) was given daily to the *GFAP/tTA; TRE/IFN-γ* mice, starting from three weeks of cuprizone exposure (W3). The corpus callosum of each mouse was collected at three weeks (W3), five weeks (W5), or eight weeks (W8) after cuprizone feeding was initiated.

### Quantitative real-time reverse transcription PCR

*GFAP/tTA;TRE/IFN-γ* mice were perfused with ice-cold phosphate-buffered saline (PBS). Total RNA was isolated from the cerebellum using the Aurum™ Total RNA Fatty and Fibrous Tissue Kit (Bio-Rad, Hercules, CA, USA). Reverse transcription was performed using the iScript™ cDNA synthesis kit (Bio-Rad). Quantitative real-time reverse transcription PCR was performed on a CFX96 RT-PCR detection system (Bio-Rad) using SYBR® Green technology. Results were, analyzed and presented as the fold induction relative to the internal control primer for the housekeeping gene *GAPDH.* The primers (5′–3′) for mouse gene sequences were as follows: *Gapdh*-f: TGTGTCCGTCGTGGATCTGA, *Gapdh*-r: TTGCTGTTGAAGTCGCAGGAG; *Ifn-γ*-f: GATATCTGGAGGAACTGGCAAAA, *Ifn-γ*-r: CTTCAAAGAGTCTGAGGTAGAAAGAGATAAT.

### Immunostaining

*GFAP/tTA; TRE/IFN-γ* mice were initially perfused with PBS only before cerebellum harvesting. The same mice were then perfused with 4% paraformaldehyde (Electron Microscopy Sciences, Hatfield, PA, USA) in PBS for 15 minutes. The brains were removed and cut coronally at approximately 1.3 mm before the bregma. The posterior parts of the brains were post-fixed overnight and embedded in O.C.T. compound (Sakura Finetek, Torrance, CA, USA). The tissue was sectioned in a series of 10 μm on a cryostat. Cryosections were treated with acetone at −20°C, then blocked with PBS containing 5% goat serum and 0.1% Triton™ X-100, and incubated overnight with the primary antibodies at 4°C. Sections were incubated with secondary antibodies for 1 h at room temperature. Coronal sections at the fornix region of the corpus callosum corresponding to Sidman sections 241-251(Sidman et al. 1971). Primary antibodies include the following anti-MBP (Abcam, ab24567, 1:700), anti-ASPA (Genetex, GTX113389, 1:500), anti-Ki67 (Abcam, AB15580, 1:100), anti-PDGFR-alpha (BD Biosciences, 558774, 1:100), and anti-Iba1(Wako Pure Chemical, 09-19741, 1:500). The fluorescent stained sections were scanned with Olympus VS-120 slide scanner and quantified by ImageJ. At least three serial sections of corpus callosum were quantified. The representative fluorescent images were acquired under Nikon A1R confocal microscope.

### Electron microscopy (EM)

For *GFAP/tTA;TRE/IFN-γ* mice, the anterior parts of the brains were immersed into EM buffer for two weeks at 4 ^o^C. EM buffer contains 4% paraformaldehyde (Electron Microscopy Sciences), 2.5% glutaraldehyde (Electron Microscopy Sciences) in 0.1 M sodium cacodylate (Electron Microscopy Sciences) at pH 7.3. The sections corresponding to the corpus callosum were trimmed, and postfixed in 1% osmium tetroxide (Electron Microscopy Sciences) in 0.1 M Sodium Cacodylate. Sections were then dehydrated in ethanol, cleared in propylene oxide, and embedded in EMBed 812 resin (Electron Microscopy Sciences). EAE mice were perfused with EM buffer, and then the lumbar spinal cords were processed, embedded, and sectioned as above. Semi-thin sections were stained with toluidine blue. Samples were next ultrathin sectioned on a Leica EM UC7 ultramicrotome. Grids were examined on a FEI Tecnai Spirit G2 transmission electron microscope. We calculated the total percentage of remyelinated axons averaged from 10 images (area = 518.3504 um^2^) in each mouse. G-ratio was calculated as the ratio of the inner diameter to the outer diameter of a myelinated axon; a minimum of 300 fibers per mouse was analyzed.

### Statistical analysis

Statistical tests were performed in Prism 8 software. No statistical methods were used to predetermine sample size. Each n value represents individual animal. All data were presented as mean ± SEM (standard error of mean). Multiple comparisons were carried out by one-way ANOVA followed by Tukey’s post hoc test; single comparisons were evaluated by unpaired t-test. Cumulative scores of EAE mice were analyzed using Kolmogorov-Smirnov method. Differences were considered statistically significant when *p* < 0.05.

## SUPPLEMENTAL INFORMATION

Figure supplement 1 is available for Figure 2.

Figure supplement 2 and 3 are available for Figure 7.

## ACKNOWLEDGEMENTS

The authors acknowledge the members of Dr. Raj Awatramani’s lab for their assistance with the VS120-S6-W slide loader system and Dr. Hongtao Chen and Mr. Lennell Reynold from the Center for Advanced Microscopy at Northwestern University for their technical support. We thank Erdong Liu for EM tissue sectioning, Dr. Vytas Bindokas from the Integrated Light Microscopy Core Facility, and Yimei Chen from the Advanced Electron Microscopy Core facility at University of Chicago for technical assistance. We also acknowledge Ani Solanki from the Animal Resource Center at University of Chicago for animal study assistance and acknowledge Sharon Way for editing the manuscript. This study was supported by NIH/NINDS R01 NS034939 (BP), the Dr. Miriam and Sheldon G. Adelson Medical Research Foundation (JRC and BP) and the Rampy MS Research Foundation (BP).

## AUTHOR CONTRIBUTIONS

Y.C. contributed to the experimental design, conducted the experiments, analyzed the data, and wrote the manuscript. R.K. and M.W. assisted with the experiments. J.C. and B.P. contributed to the experimental design and manuscript editing.

---

**Figure Supplementary 1.**
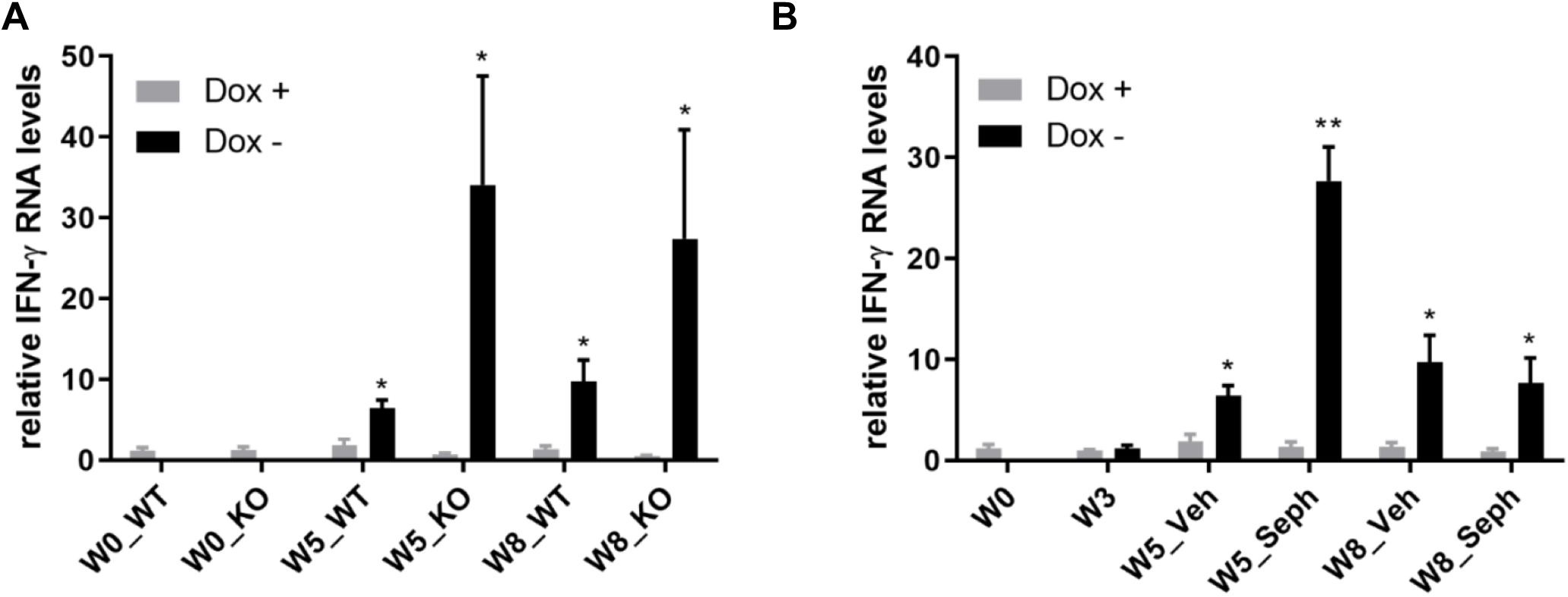
*GFAP/tTA;TRE/IFN-γ* mice express IFN-γ after release from doxycycline. Real-time qPCR analyses for detection of mRNA levels of ectopically expressed IFN-γ in the brains of *GFAP/tTA*;*TRE/IFN-γ* mice. (A) The expression of IFN-γ in the *GFAP/tTA*;*TRE/IFN-γ*/*GADD34 KO* or *WT* with the doxycycline (Dox+) and without doxycycline (Dox-). (B) The expression of IFN-γ in the *GFAP/tTA*;*TRE/IFN-γ* treated with vehicle and Sephin1 with the doxycycline (Dox+) and without doxycycline (Dox-). Data are presented as the mean ± SEM (n=4 mice/group). \**P <* 0.05, \*\**P <* 0.01. Significance based on ANOVA.

**Figure Supplementary 2.**
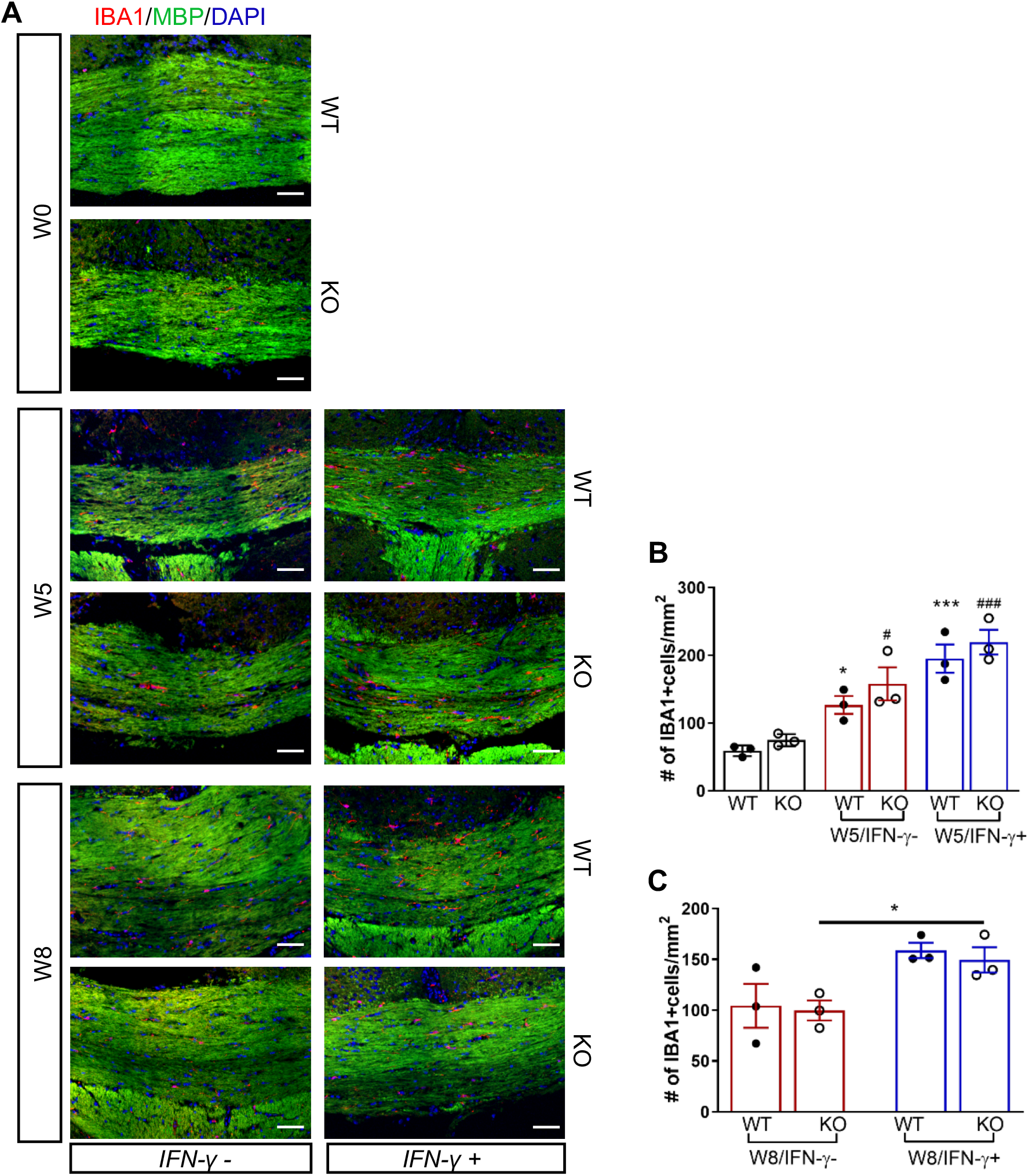
GADD34 deficiency does not affect microglial activation during remyelination in the presence of IFN-γ. The corpus callosum of *GFAP/tTA*;*TRE/IFN-γ*/*GADD34 KO* or *WT* was taken at W0, W5 and W8. (A) Immunofluorescent staining for IBA1 (a microglia marker), MBP (a myelin marker) and DAPI. Scale bar=50μm. (B) Quantification of cells positive for IBA1 in the corpus callosum areas at W0 and W5 in the absence (IFN-γ-) or presence of IFN-γ (IFN-γ+). Data are presented as the mean ± SEM (n=3 mice/group). \**P* <0.05, \*\*\**P <* 0.001 (*vs W0/WT), ^#^*P <* 0.05, ^###^*P <* 0.001 (# vs W0/KO). Significance based on ANOVA. (C) Quantification of cells positive for IBA1 in the corpus callosum areas at W8 in the absence (IFN-γ-) or presence of IFN-γ (IFN-γ+). Data are presented as the mean ± SEM (n=3 mice/group). \**P* <0.05. Significance based on ANOVA.

**Figure Supplementary 3.**
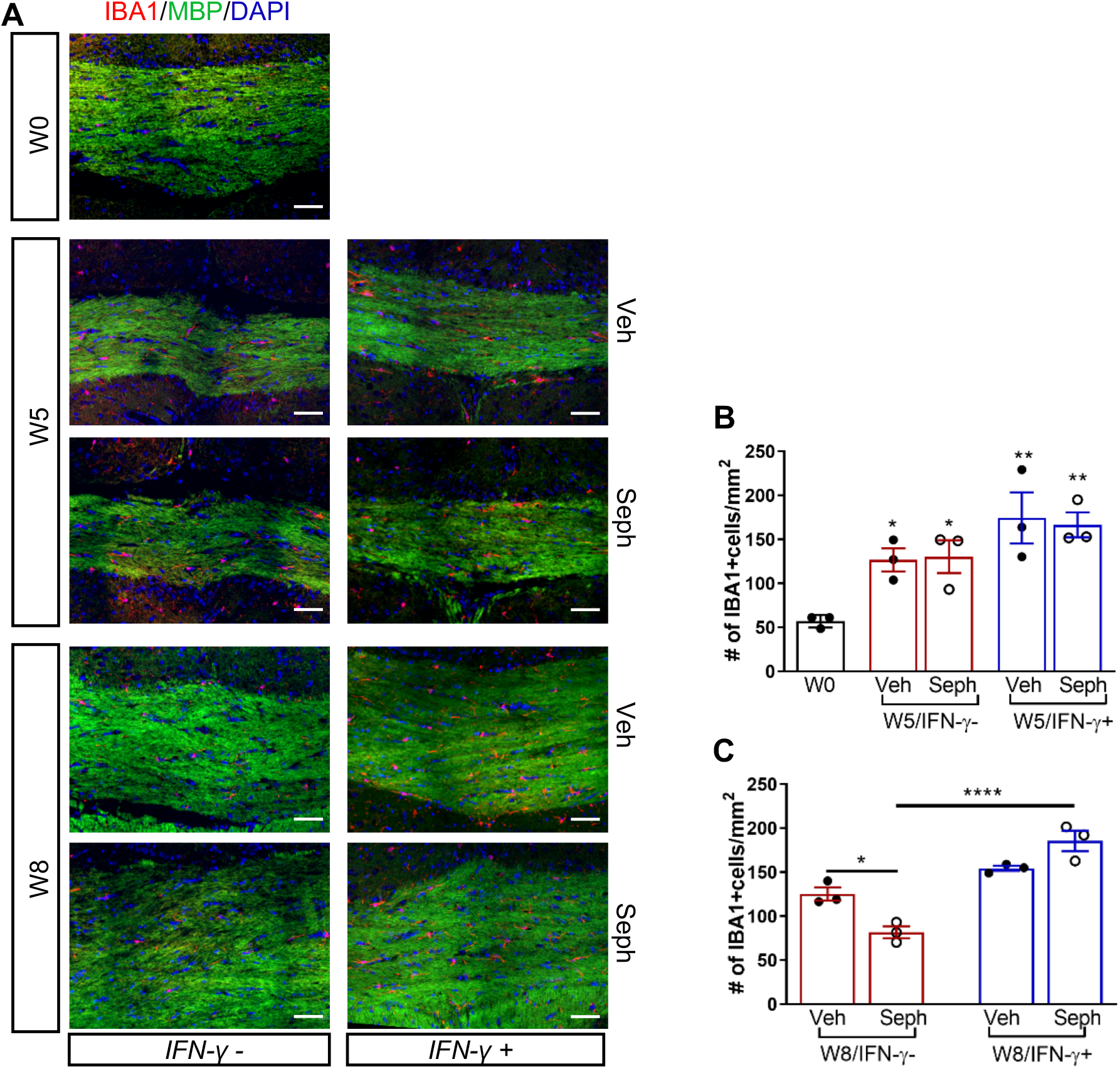
Sephin1 treatment does not affect microglial activation during remyelination in the presence of IFN-γ. The corpus callosum of *GFAP/tTA*;*TRE/IFN-γ* was taken at W0, W5 and W8 with vehicle or Sephin1 treatment. (A) Immunofluorescent staining for IBA1, MBP and DAPI. Scale bar=50μm. (B) Quantification of cells positive for IBA1 in the corpus callosum areas at W0 and W5 in the absence (IFN-γ-) or presence of IFN-γ (IFN-γ+). Data are presented as the mean ± SEM (n=3 mice/group). \**P* <0.05, \*\**P <* 0.01 (*vs W0). Significance based on ANOVA. (C) Quantification of cells positive for IBA1 in the corpus callosum areas at W8 in the absence (IFN-γ-) or presence of IFN-γ (IFN-γ+). Data are presented as the mean ± SEM (n=3 mice/group). \**P* <0.05, \*\*\*\**P* <0.0001. Significance based on ANOVA.

